# Fusion Loop Modified and Mature Dengue Virus Elicits Protective Serum with Minimal Antibody Dependent Enhancement

**DOI:** 10.64898/2026.02.18.706631

**Authors:** Yago R. Dalben, Jacob J. Adler, Rita M. Meganck, Katelyn Duenas, Lisa J. Snoderly-Foster, Longping V. Tse

**Affiliations:** Department of Molecular Microbiology and Immunology, Saint Louis University School of Medicine, St. Louis, MO, USA; Department of Molecular Microbiology, Washington University School of Medicine, St. Louis, MO, USA; MD program Saint Louis University School of Medicine, St. Louis, MO, USA

## Abstract

A balanced and safe vaccine against all four-dengue virus (DENV) serotypes is urgently needed. Although the currently licensed DENV vaccines demonstrate efficacy, they also raise concern about vaccine-associated disease enhancement, particularly in DENV-naïve individuals. The conserved, immunodominant fusion loop (FL) epitope is the target of cross-reactive, weakly neutralizing antibodies (Abs) which are associated with antibody-dependent enhancement (ADE). Previously, we developed D2-FLM, a highly mature DENV2 strain containing modification in the FL epitope making it unrecognizable to FL-Abs. Here, we grafted the FLM modification to another serotype and adapted them to replicate in Vero cells for live-attenuated vaccine (LAV) manufacturing while retaining favorable antigenic profiles, generating two new strains: D2-vFLM and D4-vFLM. Deep sequencing revealed mutations at the junction of envelope domains I and II (EDI and EDII) that appeared during adaptation of the engineered FL in mammalian cells. Importantly, both D2-vFLM and D4-vFLM showed no evidence of ADE in the presence of FL-targeting Abs. Sera from D2-vFLM immunized mice displayed strong homotypic and reduced heterotypic neutralization compared to wild-type viruses, with minimal to no ADE potential *in vitro*. Moreover, D2-vFLM immunization completely protected AG129 mice from lethal challenge with mouse-adapted D220. Collectively, these findings demonstrate that the FLM modification platform is transferable across serotypes and yields strains with favorable immunogenicity and reduced ADE risk. Our FLM approach provides a promising path toward the development of a safer tetravalent DENV LAV.

## INTRODUCTION

Dengue virus (DENV) is the most burdensome human arbovirus, infecting over 50 million people annually^1^. Despite global public health efforts, DENV-related illnesses have steadily increased over the last 75 years, with approximately half of the world’s population at risk of DENV infection^2^. Primary DENV infection often leads to asymptomatic or mild disease; however, severe cases of Dengue hemorrhagic fever/Dengue shock syndrome (DHF/DSS) can develop upon secondary infection, causing an estimated 20,000-40,000 deaths annually^1,3^. This unique etiology can be explained by antibody-dependent enhancement (ADE), in which antibodies (Abs) generated from the previous infection cross-react with but do not neutralize the incoming DENV during the next infection, amplifying the severity of the disease^4,5^. Since the first description of DENV ADE in the 1970s, our understanding of its mechanisms remains incomplete^6^. However, accumulating evidence highlights the significant role of weakly neutralizing, cross-reactive antibodies in ADE-mediated infection^7–9^. Unfortunately, there is a mismatch between our lack of understanding of ADE and the growing demographic impact of DENV infection, which is projected to escalate alongside the geographical expansion of the *Aedes* mosquito vectors and the lack of effective preventative measures.

After >20 years of development and clinical trials, three DENV LAVs are at varied phases of clinical approval, but none is recommended for DENV naive populations. The first U.S. Food and Drug Administration (FDA)-approved DENV vaccine, Dengvaxia (3 doses), is recommended for 9-16-year-old adolescents with prior DENV infection, excluding many at-risk populations. DENV naive individuals receiving Dengvaxia showed an increased risk of DENV-related hospitalization compared to placebo groups during clinical trials^10^. The manufacturer, Sanofi, has decided to halt production of Dengvaxia for the US market in 2026. Qdenga (2 doses), a vaccine approved by the European Medicines Agency, voluntarily withdrew its application for FDA approval and, due to low incidence, is unable to provide key clinical trial data addressing vaccine safety concerns in naive individuals infected with DENV3 and DENV4^11,12^. The final candidate in the pipeline, TV005 (single dose), elicits protection against DENV1 and DENV2 in a 2 years follow-up clinical trials^13^. However, conceptually, it is constrained by the same philosophy and technical limitations as its predecessors. It elicits Abs targeting all epitopes, raising similar safety concerns about ADE-prone Abs at later time (> 5 years) after vaccination. Twenty years later, our understanding of ADE has advanced, emphasizing the need to incorporate strategies to minimize ADE-prone Abs into the development of next-generation DENV vaccines to ensure safety and efficacy in naive populations.

The pre-membrane (prM) protein and fusion loop (FL) are epitopes known to be targets of weakly neutralizing, ADE-prone antibodies. PrM shields the FL peptide from premature fusion during egress, while the FL mediates viral membrane fusion with the endosome during entry. The essential functions of prM and FL make them challenging to mutate. Indeed, DENV prM shares approximately 66-80% sequence identity among the four serotypes, while the core 13-amino acid FL is 100% conserved in mosquito-borne orthoflaviviruses. Although prM is normally released after egress, many DENV particles remain partially immature and retain high levels of prM. Methods have been developed to generate mature virions by overexpressing furin protease in producer cells; however, for LAVs, the progeny virus produced in the human body will revert to the immature phenotype. Therefore, current DENV-LAVs present both prM and FL epitopes and elicit strong but potentially undesired antibody responses against them. Supporting this, sera from primary human DENV infections show that a significant proportion of Abs target prM and FL and are weakly neutralizing and ADE-prone, which poses substantial ADE risk for subsequent infections^14^. This issue has been well recognized in vaccine development, and efforts to mask, mutate, or remove prM and FL epitopes have been applied in subunit and mRNA-LNP vaccines^15^. However, such protein-engineering strategies cannot be directly translated to LAVs, where these elements are required for viral function.

The highly effective flavivirus vaccine against Yellow Fever virus, Y17D^16^, as well as all licensed DENV vaccines are LAVs^17^. LAVs mimic natural infection and elicit strong humoral and cell-mediated responses against both structural and non-structural proteins. Importantly, unlike other vaccine platforms which requires multiple boosters, TV005 (the most advanced DENV-LAV) provides protection with a single dose^18^, a critical feature for low-income regions where multi-dose regimens are difficult and where partial immunity between doses may increase the risk of ADE.

Previously, our group reported genetically modified mature DENV and a FL-mutated, fully mature DENV2 (D2-FLM) that is insensitive to FL- and prM-targeting Abs^19^. In this study, we extended this strategy by grafting the FLM mutations onto a genetically distinct serotype (DENV4) and adapting both FLM viruses in Vero cells to generate D2- and D4-vFLM, suitable for large-scale LAV manufacturing. Both vFLM strains retained their intended antigenic properties and did not cause ADE in the presence of FL-Abs. Immunization with D2-vFLM elicited robust neutralizing antibody responses against the homotypic virus and, to a lesser extent, the heterotypic virus. Importantly, immune sera from D2-vFLM viruses show minimal ADE activity *in vitro*. Furthermore, AG129 mice immunized with D2-vFLM were fully protected against lethal challenge with mouse-adapted D220. Together, these findings suggest that the FLM mutations represent a promising and potentially safer LAV platform, capable of generating favorable humoral responses to protect DENV-naïve individuals while minimizing vaccine-related ADE.

## RESULTS

### Generation and characterization of Vero-adapted D2-vFLM

All currently licensed or advanced DENV LAVs are manufactured in Vero cells^20,21^. Thus, the ability to replicate in this cell line is a critical requirement for further development of the FLM series as a potential LAV candidate. As we previously reported, D2-FLM strain replicates to levels comparable to isogenic DV2 in C6/36 cells^19^. However, D2-FLM is attenuated in Vero cells; in multi-cycle growth curves, no detectable titer was measured in over five days. To understand the growth defect, we performed immunostaining with anti-NS3 antibody at 24hpi. Robust NS3 staining was detected in DV2 infected cells, but no signal was observed in D2-FLM infected cells, suggesting that the block occurs early in infection, prior to NS3 translation **(Fig. 1A)**.

**Figure 1.**
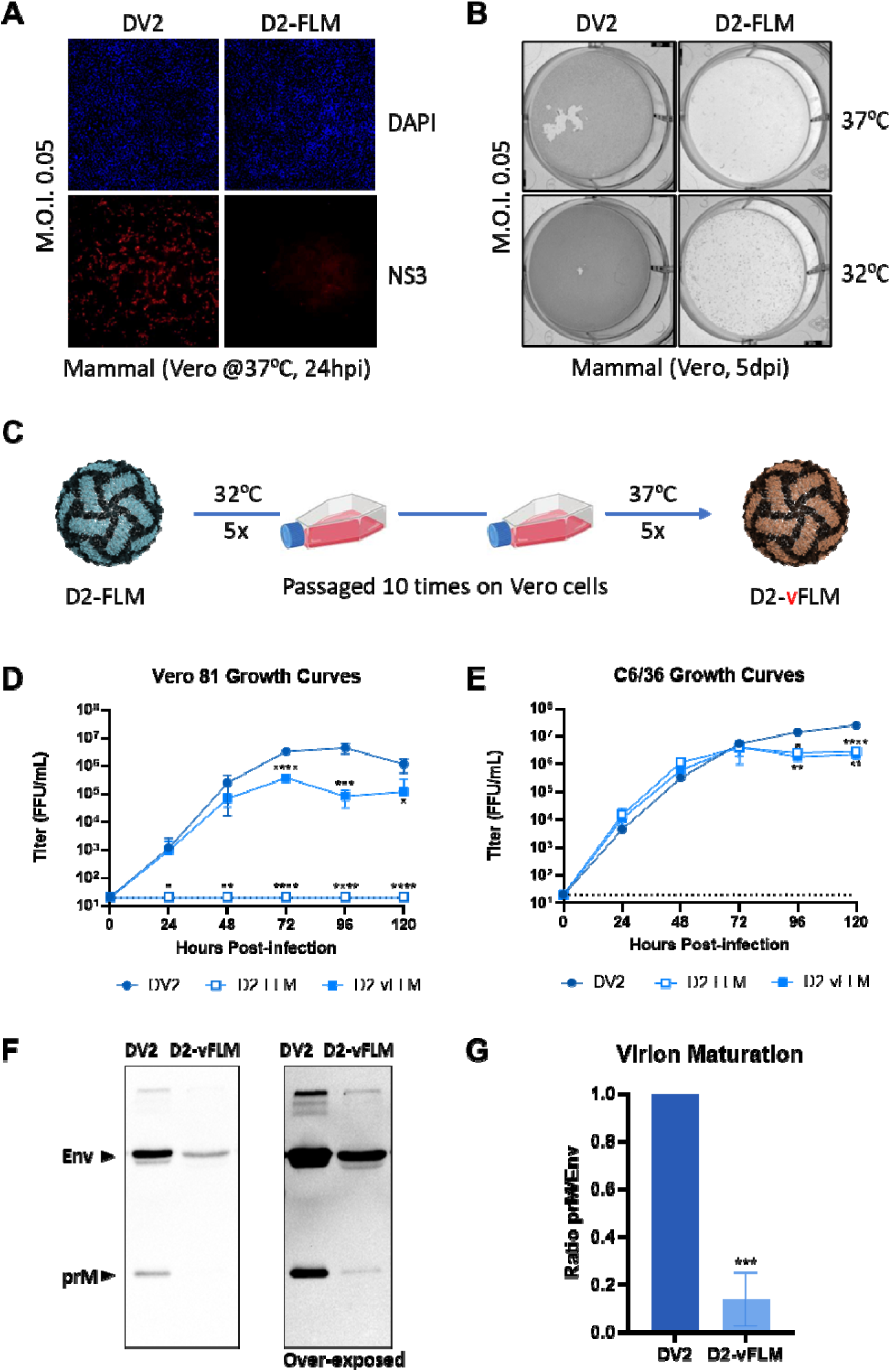
Adaptation and characterization of D2-vFLM D2-FLM. **A)** Immunofluorescent staining of DV2 and D2-FLM infected Vero-81 cells with a-NS3 antibody at 24 hours post-infection. **B)** Combined staining of prM and E of DV-2 and D2-FLM infected Vero-81 cells at 32°C and 37°C at 5 days post-infection. **C)** Schematic of D2-vFLM mammalian cell adaptation. Multiple-step growth curves of DV2, D2-FLM, and D2-vFLM in **D)** Vero-81 and **E)** C6/36 cells. **F)** Western blot of DV2 and D2-vFLM with a-E and a-prM antibodies. Right: over-exposure to visualize prM in D2-vFLM. **G)** Quantification of prM:E ratio of D2-vFLM, normalized to DV2. Growth curves were analyzed by two-way ANOVA with multiple comparison to DV2 control at different time points. Western blot quantification was analyzed by Students *t-*test.

Because C6/36 cells are cultured at 32°C whereas Vero cells are maintained at 37°C, we hypothesized that D2-FLM might be temperature sensitive. To test this, Vero cells were infected with DV2 or D2-FLM (MOI = 0.05) and incubated for five days at either 32°C or 37°C. Infectivity was assessed by immunostaining for prM and E proteins at day 5 post-infection. DV2 showed robust staining at both temperatures, with nearly complete infection across the well. In contrast, D2-FLM exhibited moderate staining at 32°C with scattered foci but showed no evidence of replication at 37°C **(Fig. 1B)**. These findings indicate that D2-FLM is partially temperature sensitive and that 32°C is permissive for low-level replication in Vero cells, allowing for experimental tissue-culture adaptation. To achieve this, D2-FLM was serially passaged on Vero cells at 32°C for 3–5 passages to promote mammalian cell adaptation, followed by an additional 3–5 passages at 37°C for temperature adaptation **(Fig. 1C)**. Infectivity, assessed by prM and E staining, increased progressively with passage number (data not shown). After the final passage, we collected the adapted virus and designated it D2-vFLM, where “v” indicates Vero-adapted.

D2-vFLM replicates in Vero cells at 37°C, reaching peak titers of ∼4× 10^5^ FFU/mL at day 3 post infection **(Fig. 1D)**. Notably, D2-vFLM remained mildly attenuated, with a 10-fold reduction in peak titer compared with DV2 (∼3×10^6^ FFU/mL; **Fig. 1D)**. In C6/36 cells, D2-vFLM grows with similar kinetics to D2-FLM, reaching ∼5×10^6^ FFU/ml at 3dpi, while DV2 reaches peak titers of 1×10^7^ FFU/mL at 5dpi **(Fig. 1E)**. The maturation phenotype of D2-FLM is generated by the genetically modified furin cleavage site in prM^22^. As expected, western blot analysis confirmed that D2-vFLM retains the highly mature virion phenotype **(Fig. 1F)** with an 80% reduction of unprocessed prM compared to DV2 in normal Vero-81 producer cells **(Fig. 1G)**.

To assess virion stability, we performed an infectivity assay using nano-luciferase (nLuc) reporter DENV. D2-vFLM-nLuc was rederived by reverse genetics and incubated at temperatures from 4°C to 55°C and pH levels from 5.4 to 8.4. Infectivity was normalized to untreated controls and compared with DV2. Interestingly, DV2 displayed up to a 2.5-fold increase in infectivity after incubation at temperatures from 32-42°C. In contrast, D2-vFLM infectivity was stable from 4-37°C, with a noticeable decrease beginning at 42°C. As expected, treatment at 55°C completely inactivated both D2-vFLM and DV2 **(Fig. S1A)**. Although some variation was observed between experiments, D2-vFLM displayed similar pH stability to DV2 from pH 5.4 to pH 8.4. At acidic pH, a 20–40% loss of infectivity was observed for both viruses **(Fig. S1B)**. These results suggest that the thermostability and pH stability of D2-vFLM are similar to wild-type viruses.

#### Mutations in the structural protein are selected for during Vero adaptation

To confirm the stability of FLM modifications and to identify compensatory mutations after Vero adaptation, we deep sequenced the D2-vFLM virus. First, the FLM mutations are stable and do not revert to the wild-type sequence after eight passages. Second, we identified a total of six fixed mutations in both the structural and non-structural genes, including prM-E18D, Env-P53T, Env-T171A, Env-K204R, NS2A-L181F, and NS4b-L112F **(Table 1)**. To determine which mutations were driving adaptation, we generated two variants: D2-vFLM-structural and D2-vFLM-nonstructural, which carried only the structural or non-structural adaptation mutations, respectively. Individual genome fragments containing the desired mutations were cloned, and infectious clones were recovered in C6/36 cells using reverse genetics **(Fig. 2A)**. Working stocks were subsequently prepared by propagation in either C6/36 cells (insect-grown) or Vero cells (mammalian-grown). Deep sequencing of all stocks confirmed the presence of the core FLM mutations and the intended adaptation mutations **(Table 1)**.

**Figure 2.**
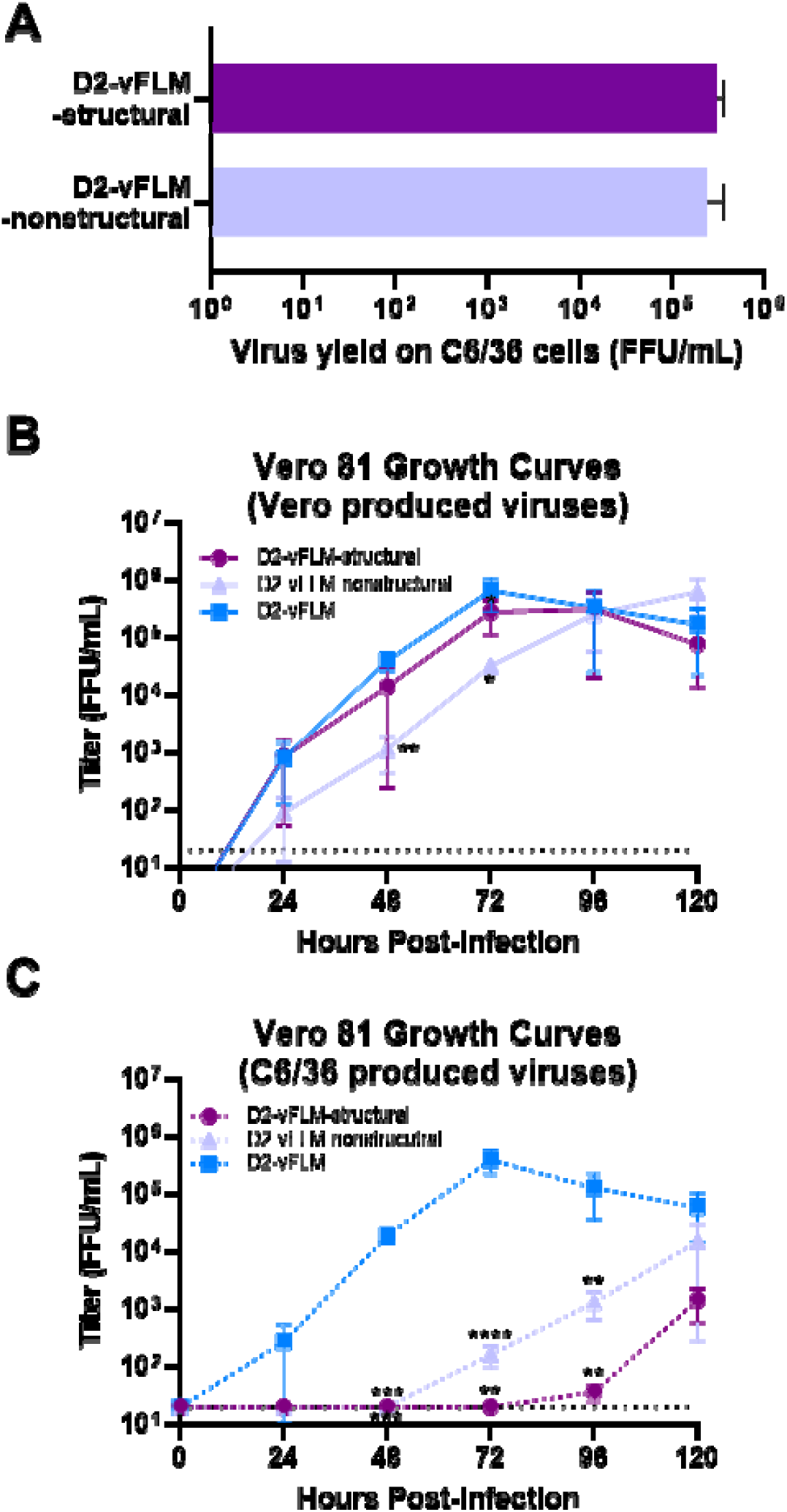
Compensatory mutations in structural proteins are important for Vero-81 adaptation. **A)** Virus yield of D2-vFLM-strucutral and -nonstructural infectious clones grown on C6/36 cells. Multiple-cycle growth curves of **B)** Vero grown or **C)** C6/36 grown virus D2-vFLM-strucutral and -nonstructural viruses on Vero-81 cells. Growth curves were analyzed by two-way ANOVA with multiple comparison to D2-vFLM control at different time points.

**Table 1.** Engineered and acquired mutations for different D2-vFLM mutants in Vero-81 cells.

|  | D2-vFLM | D2-vFLM-structural | D2-vFLM-nonstructural |
| --- | --- | --- | --- |
| **Structural mutations** |  |  |  |
| prM-E18D | 100% | 100% (I) | - |
| prM-L41V | - | - | 100% (A) |
| Env-P53T | 100% | 100% (I) | - |
| Env-T171A | 100% | 100% (I) | 95.4% (A) |
| Env-K204R | 100% | 100% (I) | - |
| Env-T404K | - | - | 97.3% (A) |
| **Nonstructural mutations** |  |  |  |
| NS2A-L181F | 100% | - | 100% (I) |
| N24b-L112F | 100% | - | 100% (I) |
| NS1-M89V | - | 31% (A) | - |
| NS3-D409E | - | 14% (A) | - |
| NS4b-V115A | - | 13% (A) | - |

(I): Intended mutations; (A): *Acquired mutations*; - Not Applicable

For D2-vFLM-structural, the mammalian-grown stock contained three spontaneous mutations at low frequency: NS1-M89V (31%), NS3-D409E (14%) and NS4b-V115A (13%) **(Table 1)**. In contrast, sequencing of the mammalian-grown D2-vFLM-nonstructural stock revealed three structural mutations at >95% frequency: prM-L41V, Env-T171A (present in the original D2-vFLM), and T404K, yielding a virus with the intended non-structural changes plus three additional structural changes **(Table 1)**. These findings suggest that structural protein mutations are strongly favored to co-evolve with FLM mutations in mammalian cells.

To assess replication kinetics, we performed multi-cycle growth curve experiments with mammalian-grown D2-vFLM-structural and D2-vFLM-nonstructural viruses in Vero cells. D2-vFLM-structural exhibited growth kinetics similar to D2-vFLM, reaching peak titers of ∼3×10^5^ FFU/mL at 3 dpi. In contrast, replication of D2-vFLM-nonstructural was slower, peaking at 5 dpi **(Fig. 2B)**. Interestingly, when insect-grown virus stocks were used for Vero-81 growth curves, both D2-vFLM-structural and -nonstructural viruses showed a 48h delay in replication, and viral titer did not exceed 10^5^ FFU/mL after 5 days **(Fig. 2C)**. These data indicate that C6/36-grown D2-vFLM displays non-genetic attenuation in mammalian cells, providing an added safety feature of the LAV by reducing the risk of insect-to-human transmission.

### FLM mutations can be directly grafted onto DENV-4

We envision the FLM mutations as platform technology for developing a tetravalent DENV LAV. As a proof-of-concept, we examined the compatibility of the FLM mutations in the most genetically distinct serotype, DENV4. The core FLM mutations were introduced into a DENV4-Sri Lanka 1992 backbone and a viable virus was successfully recovered in C6/36 cells using reverse genetics. Multi-cycle growth curve analysis of D4-FLM on Vero cells revealed a strong attenuation phenotype like D2-FLM. D4-FLM showed a >2-log reduction in peak titer (∼10^4^ FFU/mL) compared to the isogenic DV4 control (∼10^6^-10^7^ FFU/mL) at 4dpi **(Fig. 3A)**. These findings indicate that FLM mutations are transferable to other DENV serotypes and have a predictable attenuation on Vero cell replication.

**Figure 3.**
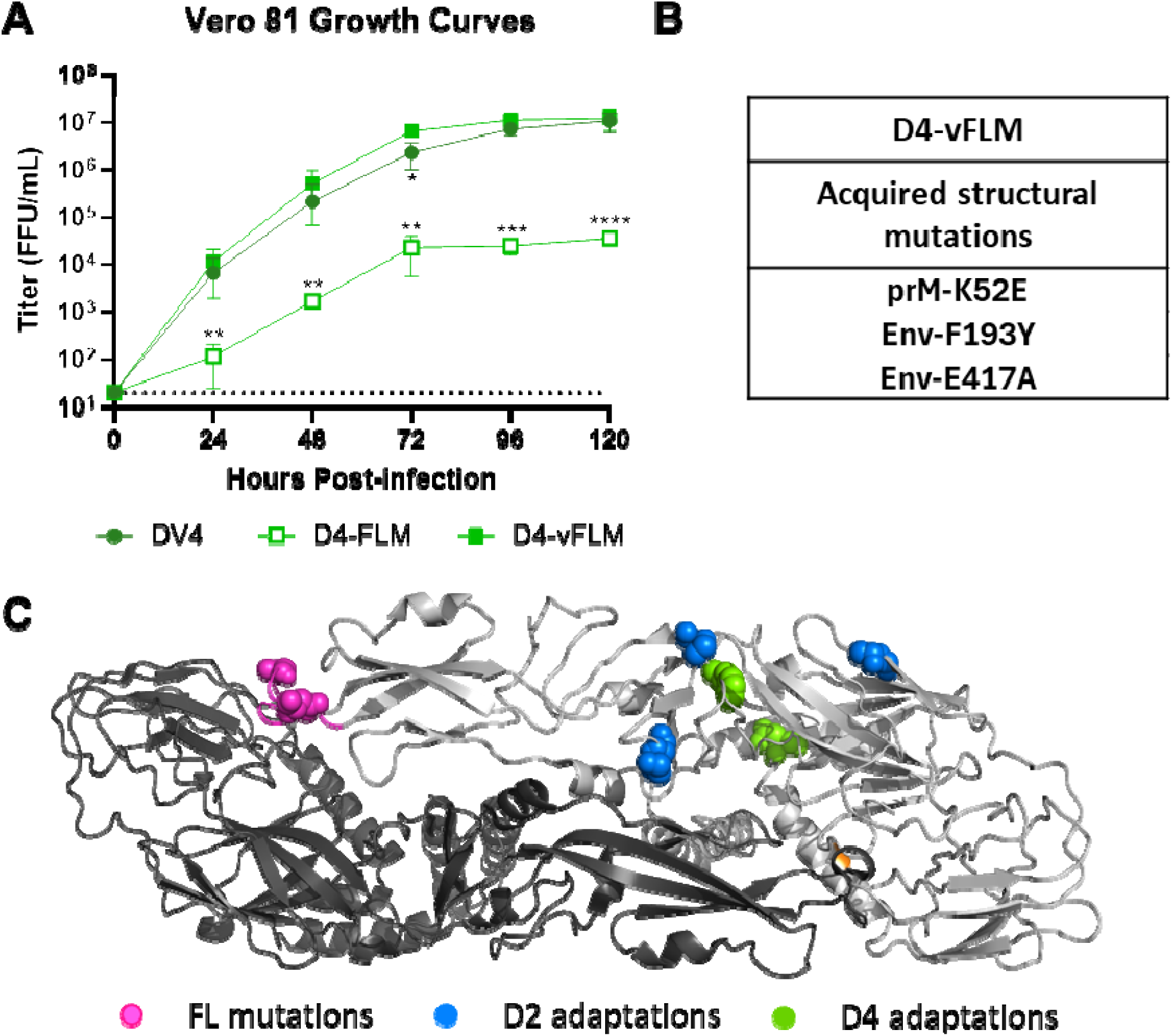
Grafting and adaptation of FLM mutations on DENV4. **A)** Growth curves of DV4, D4-FLM and D4-vFLM on Vero-81 cells. **B)** Acquired mutations of D4-vFLM after serial passage in Vero-81 cells. **C)** Structural model of DENV-Envelope-dimer with color coded FL mutations and Vero adaptation mutations from DENV2 and DENV4. Growth curves were analyzed by two-way ANOVA with multiple comparison to DV4 control at different time points.

To overcome this growth defect, we subjected D4-FLM to serial passage in Vero cells at 37°C for four rounds, similar to the approach used with D2-FLM. The adapted virus, termed D4-vFLM, regained replication capacity comparable to wild-type DENV4, reaching titers of ∼10^6^-10^7^ FFU/mL by d4pi **(Fig. 3A)**. Deep sequencing confirmed retention of the FLM mutations, along with additional adaptive mutations in structural proteins: prM-K52E (97.4%), Env-F193Y (97.5%), and Env-E417A (100%). No mutations were identified in non-structural proteins **(Fig. 3B)**. Although no overlapping adaptive mutations were identified between D2-vFLM and D4-vFLM, structural adaptations were concentrated at the EDI and EDII junction of the envelope protein **(Fig. 3C)**, suggesting a possible structural hotspot for FL mutations compensation.

#### Antigenic properties of D2- and D4-vFLM

We previously demonstrated that D2-FLM exhibits distinct antigenic properties, becoming insensitive to FL-targeting antibodies (FL-Abs). However, there was a possibility that additional structural mutations in D2-vFLM might alter these unique antigenic features. To address this, we performed foci reduction neutralization titer (FRNT) assays on Vero cells testing D2-vFLM and D4-vFLM against a panel of monoclonal antibodies. Consistent with earlier findings, FRNT_50_ values for non-FL antibodies, including prM (2H2, non-neutralizing), EDE antibodies (EDE1-C10 and EDE2-B7), EDIII antibody (2D22), and EDI antibody (3F9), showed no significant differences between D2-vFLM and DV2. In contrast, D2-vFLM displayed varying degrees of reduced sensitivity to FL-Abs (1M7, 1L6, and 4G2) compared to DV2 **(Fig. 4B)**. Similarly, D4-vFLM exhibited altered antigenicity, with selective loss of sensitivity to FL-Abs (1M7, 1L6, and 4G2) while remaining minimally affected by broadly neutralizing EDE antibodies (EDE1-C10 and EDE2-B7) **(Fig. 4B)**.

**Figure 4.**
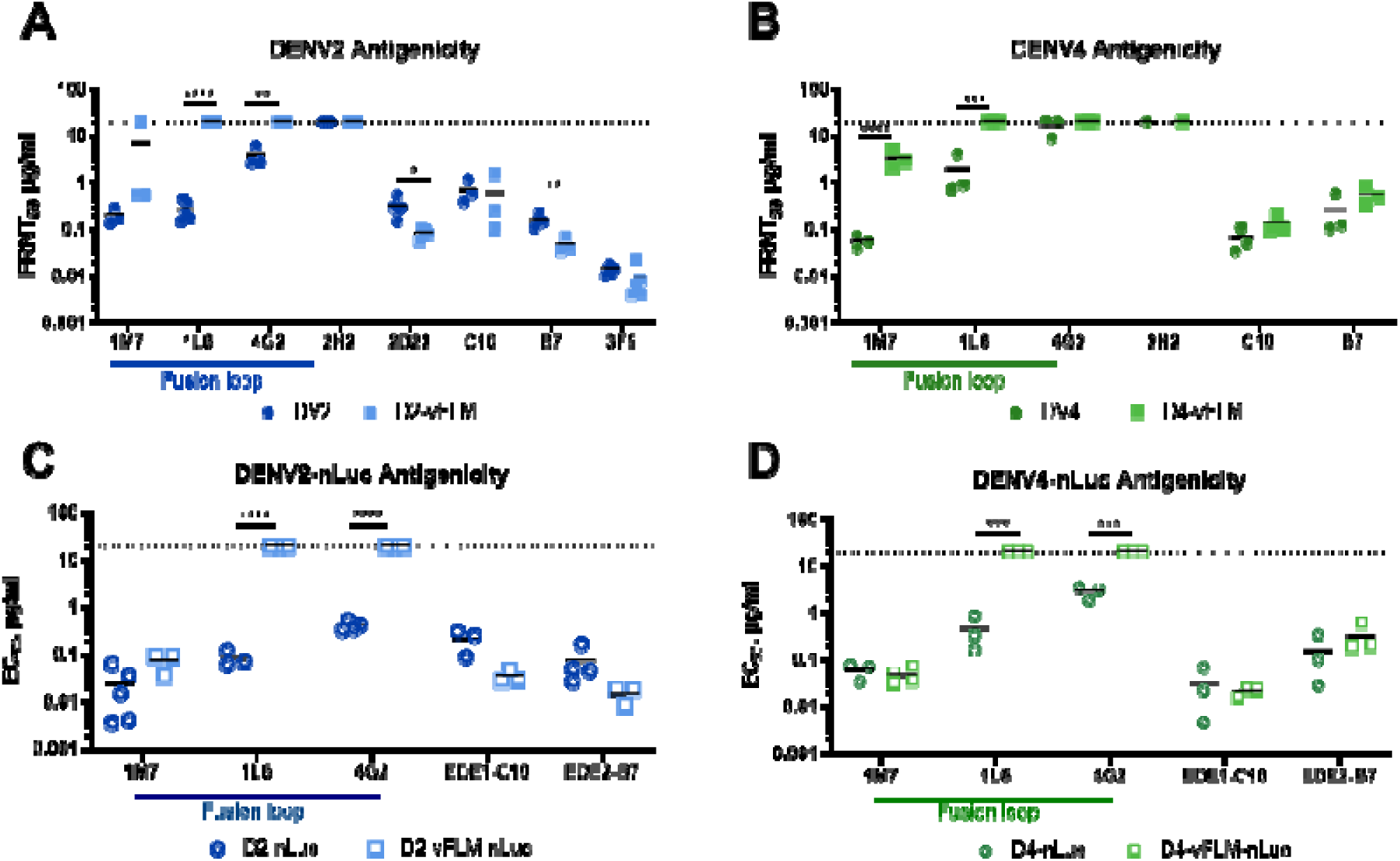
Antigenicity of DENV and DENV-vFLM. FRNT_50_ of **(A)** DV2 and D2-vFLM or **(B)** DV4 and D4-vFLM against mAbs on Vero-81 cells. EC_50_ of **(C)** DV2-nLuc and D2-vFLM-nLuc or **(D)** DV4-nLuc and D4-vFLM-nLuc against mAbs on Vero-81 cells. Statistics were performed by Students *t-*test.

To facilitate future studies, we generated D2-vFLM-nLuc and D4-vFLM-nLuc reporter viruses using reverse genetics based on the design reported previously. Neutralization assays with these reporter viruses yielded EC_50_ values that were consistently ∼10-fold lower (more sensitive) than FRNT_50_ values across multiple antibodies **(Fig. 4C and 4D)**, though the relative potency remained largely unchanged. One notable exception was observed with D4-vFLM-nLuc against 1M7: the nLuc EC_50_ (∼0.05 μg/ml) was comparable to DV4, whereas the FRNT_50_ of D4-vFLM was significantly higher (∼2.78 μg/ml) **(Fig. 4D)**. This discrepancy is not readily explained; however, deep sequencing revealed a unique mutation (prM-E89S) in D4-vFLM-nLuc, while D4-vFLM carried prM-K52E. These differences may provide testable hypotheses for future investigation but are beyond the scope of the present study.

### Immunization with D2-vFLM generates neutralizing serum comparable to DV2

To compare the immunogenicity of the D2-vFLM virus with wild-type DV2, we immunized 3-4-month-old C57BL/6 mice (both male and female) via intramuscular (I.M.) injection in the hind leg with 2.5×10^5^ FFU of DV2 (n=10) or D2-vFLM (n=11) adjuvanted with AddaVax. A relatively high dose was chosen compared with the human (human dosage: 10^3^-10^5^ PFU per dose) due to the inability of DENV to replicate efficiently in immunocompetent mice. Mice received three immunizations as prime/1°boost/2°boost administered at two-week intervals. Homotypic and heterotypic serum neutralization titers were measured using our nLuc-based neutralization assay. Pooled serum from pre-immunization, after prime, after 1°boost, and after 2°boost shows steady increase in neutralization titer **(Fig. 5A**; individual neutralization curves in **Fig. S2)**. Robust seroconversion started after 1°boost and peaked after 2°boost. Homotypic EC_50_ against DENV2 reaches ∼1:1000 for both viruses. Heterotypic EC_50_ against DENV4 were minimal and only appears after 2°boost.

**Figure 5.**
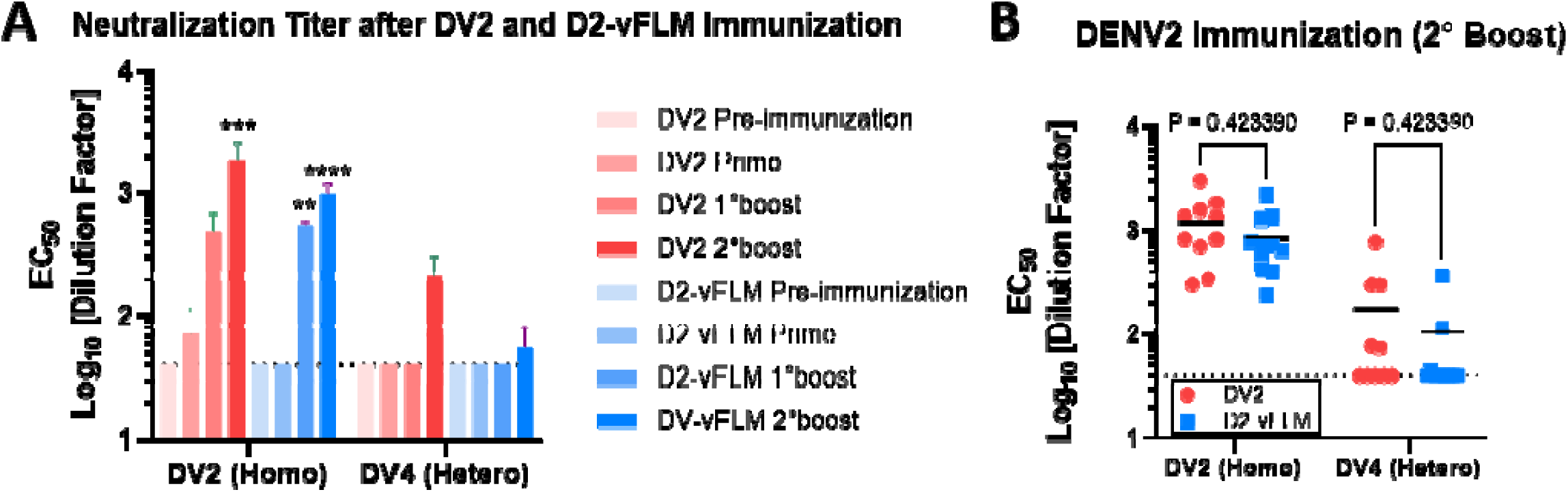
Immunogenicity of DVs and D2-vFLMs *in vivo*. **A)** Homotypic (against DV2-nLuc) and heterotypic (against DV4-nLuc) EC_50_ of pooled serum of DV2 and D2-vFLM from pre-immunization, 2wk after prime, 2wk after 1°boost, and 2 wk after 2°boost. **B)** Homotypic and heterotypic EC_50_ of individual DV2 and D2-vFLM serum from immunized C57BL/6 mice after 2°Boost. EC_50_ of pooled serum were analyzed by two-way ANOVA with multiple comparisons compared to pre-immunization. Mean EC_50_ of the 2°boost was compared by Students *t-*test.

To gain better understanding of seroconversion of individual animals, we perform neutralization assay on each serum after 2°boost. EC_50_ valves span over a 10-fold range, from 1:300 to 1:3000 while the mean EC_50_ values against homotypic DENV2 were ∼1:1000 for DV2 and 1:800 for D2-vFLM; against the heterotypic DENV4, mean EC_50_ values were 1:180 and 1:110, respectively **(Fig. 5B**; individual neutralization curves in **Fig. S3A and S3B)**. Our data suggests that the FLM mutations in D2-vFLM may modestly reduce immunogenicity, potentially due to the loss of immunodominant epitopes, although this difference did not reach statistical significance. Overall, D2-vFLM viruses elicited neutralizing antibody responses comparable to those induced by DV2.

### Serum from D2-vFLM shows minimal to no ADE responses against heterotypic viruses

To compare the heterotypic ADE potency of D2-vFLM and DV2 serum, we developed a high-throughput nLuc-based ADE assay. In our assay, we evaluated three antibodies targeting distinct antigenic regions, including 4G2 (FL-Ab), 2H2 (prM-Ab), and B7 (EDE-Ab), using DV4-nLuc reporter viruses on K562 cells. As expected, all three antibodies enhanced DV4-nLuc infection at peak dilution/concentrations of 1:100 (4G2 hybridoma) **(Fig. 6A)**, 1:100 (2H2 hybridoma) **(Fig. 6B)**, and 1 µg/ml (B7) **(Fig. 6C)**. In contrast, 4G2 did not enhance infectivity of D4-vFLM, while 2H2 induced only a modest level of enhancement. Because the EDE epitope is preserved in the vFLM viruses, D4-vFLM exhibited similar peak ADE concentrations for B7 compared to DV4 **(Fig. 6C)**. We benchmarked our results against published data obtained using conventional flow cytometry-based ADE assays and flaviviral single-round infectious particle (SRIP) systems. Results from published ADE assays can be variable; however, our results fall within the reported range in both fold-increase (10–100-fold) and antibody concentration (0.1–5 µg/ml 4G2-Ab) for enhancement^23–25^. These data validate our nLuc-based ADE assay, and that the FLM mutations render DENV inert to ADE mediated by FL-Abs.

**Figure 6.**
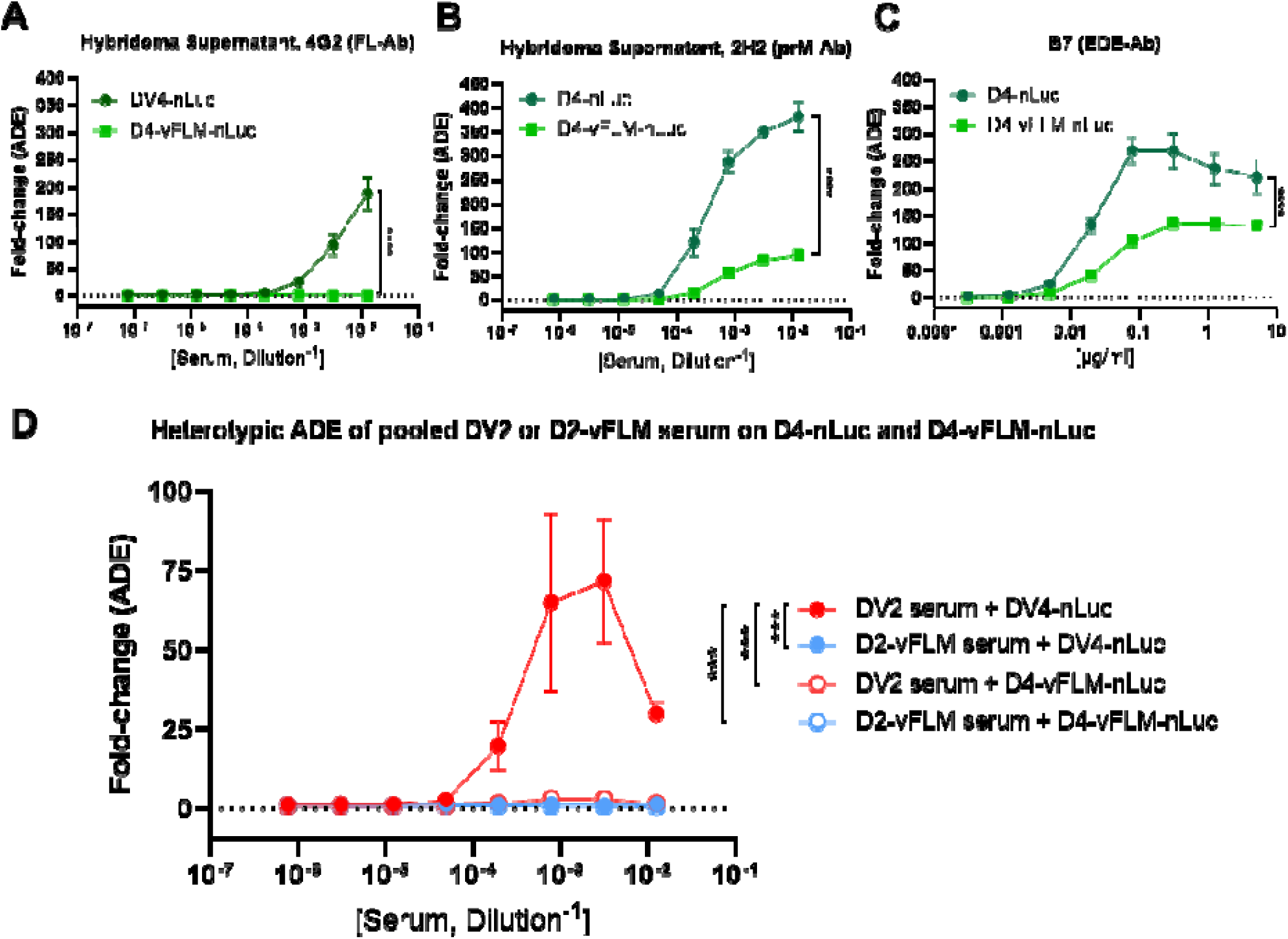
High-throughput nLuc-based ADE assay shows no heterotypic ADE from serum isolated from D2-vFLM immunized mice. Monoclonal Ab control and fold-increase in infectivity of DV4-nLuc and D4-vFLM-nLuc against **A)** 4G2, **B)** 2H2, and **C)** B7 in K562 cells. **D)** Heterotypic ADE against pooled serum from DV2 and D2-vFLM immunized mice that are positive against DV4. Error bars denote ±1 S.D. Data was analyzed by two-way ANOVA; statistical significance shown corresponds to the interaction between groups.

Based on the neutralization assay, we analyze the ADE potency of 2°boost serum of DV2 (n=5) and D2-vFLM (n=4) that are positive against heterotypic DV4. Pooled, positive sera from DV2-immunized mice against the heterotypic DV4-nLuc exhibited approximately 75-fold increase at peak compared to no serum control **(Fig. 6D)**. In contrast, pooled, positive D2-vFLM-immunized sera exhibited negligible ADE. Additionally, DV2-immunized sera did not enhance D4-vFLM-nLuc infection, further support that the enhancing Abs in the polyclonal serum target FL- and prM-epitopes **(Fig. 6D)**. Overall, our data demonstrates that FLM modifications achieve a key immunological objective by minimizing heterotypic ADE. Full neutralization and ADE curves of individual serum samples can be found in supplementary figure 4 **(Fig. S4)**.

### D2-vFLM immunization protects mice from lethal challenge in AG129 mice

To test *in vivo* protection, male and female AG129 mice (4-6-months-old) were immunized with DV2 (n=4, one animal died before challenge); D2-vFLM (n=5); or mock immunized (n=6, one animal died before challenge) with the same dosing regimen as above. Three weeks after the final boost, mice were challenged with 5×10^5^ FFU of mouse-adapted D220 via tail-vein intravenous (I.V.) injection following published protocols^26^. In mock immunized mice, weight loss was observed beginning at 2dpi, and lethality at 5dpi, reaching 83% mortality by 10dpi **(Fig. 7A and 7B)**. Viral titers were detected in sera from mock immunized mice, peaking at 6.2×10^6^ FFU/ml on d3pi **(Fig. 7C)**. D2-vFLM immunized mice showed complete protection from weight loss, viral titer (limit of detection: 200 FFU/ml), and death, similar to DV2 immunized mice **(Fig. 7)**. Our data demonstrates D2-vFLM immunization can protect from homotypic infection *in vivo*.

**Figure 7.**
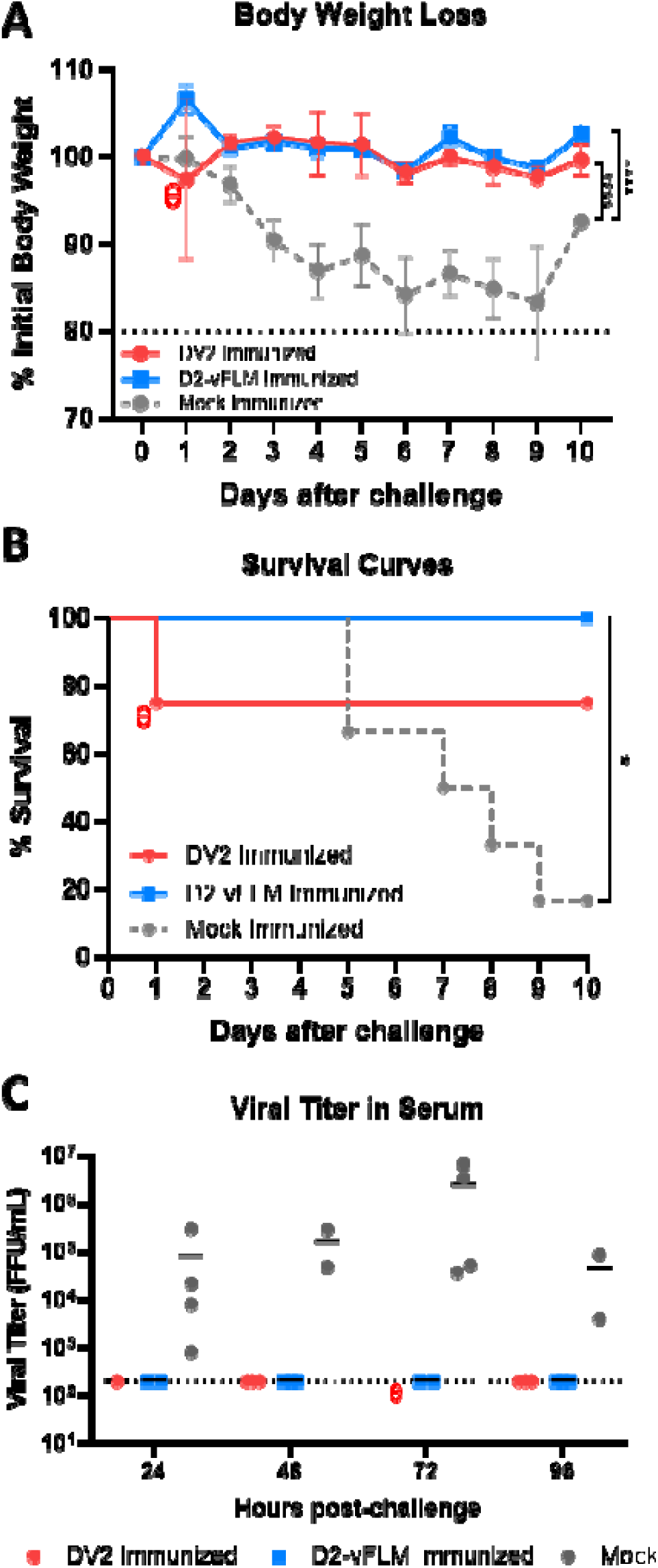
D2-vFLM immunization protects AG129 mice from lethal challenge. AG129 mice were immunized with DV2 (n=4), D2-vFLM (n=5), or mock (n=6) immunized and challenged with D220 virus. **A)** Percent weight loss, analyzed by linear mixed-effects model with repeated measures. Pairwise group comparisons were performed using estimated marginal means with Tukey adjustment. **B)** Survival curves up to 10 days post challenge, assessed with Log-rank (Mantel–Cox) test, and pairwise comparisons performed with Bonferroni correction for multiple testing. **C)** Serum viral titer from d1pi - d4pi. Individual mice are only bled twice per week for humane treatment. *P* < 0.05 was considered statistically significant. Θ represents unexpected mouse death on d1pi in the DV2 immunization group.

## DISCUSSION

Antibodies targeting the FL- and prM-epitopes have been associated with ADE^8,9^. Therefore, Dengue and flavivirus vaccine design strategies often aim to mask or eliminate the FL epitopes^15,27–29^. However, the requirement for a fully functional FL in the LAV platform has posed a significant challenge for the development of a FL-modified DENV-LAV^30^. Previously, using directed evolution, we introduced mutations in the FL and showed that the resulting virus, D2-FLM, exhibited the intended antigenic property of reduced recognition by FL-targeting antibodies^19^. However, these mutations also resulted in the inability of D2-FLM to grow in Vero-81 cells, the production cells for DENV-LAV manufacturing. In this study, we extended our previous work by adapting the FLM virus to enable growth in Vero cells and generating two Vero-adapted FLM viruses, D2-vFLM and D4-vFLM, and evaluating D2-vFLM potential as LAV candidates.

D2- and D4-vFLM both grow in Vero-81 cells, supporting FLM modifications can be transfer to other DENV serotypes as well as orthoflaviviruses for direct downstream vaccine manufacturing. Like the parental FLM viruses, vFLM strains have lost or reduced neutralization and enhancement by FL-targeting antibodies. Mice immunized with D2-vFLM seroconverted and showed neutralization potency comparable to their WT counterparts. Although we did not compare the cell-mediated responses between D2-vFLM and DV2, we expect the mutations of the vFLM virus would have minimal effect on T-cell epitopes as most of the CD4 and CD8 epitopes are from non-structural proteins, such as NS3^31,32^. Importantly, using an *in vitro* ADE assay, our data indicates that D2-vFLM induces qualitatively different antibody responses that do *not* enhance heterotypic infection. *In vivo* protection was evaluated in a homotypic format and D2-vFLM immunized AG129 mice were completely protected from mouse-adapted D220 challenge. Overall, DENV strains carrying FLM modifications elicited a safer antibody profile while maintaining immunogenicity in mice, a critical step to improve the safety of DENV-LAV.

Because the FLM mutations are deleterious to viral replication in Vero-81 cells, compensatory mutations were required to restore fitness. We did not identify consensus compensatory mutations between different serotypes, but they are primarily located in the hinge region between EDI and EDII. A plausible hypothesis is that the FLM modifications may influence the conformational transition between the pre-fusion dimeric and post-fusion trimeric forms that are only affected in mammalian cells^33,34^. We are currently investigating this phenomenon to identify universal compensatory mutations that could eliminate the need for adaptation of additional strains.

The generation of FLM viruses not only holds potential for incorporation into next-generation DENV-LAVs to improve safety but also provides valuable tools for characterizing anti-DENV sera. Owing to their unique antigenicity, which reduces recognition by FL- and prM-targeting antibodies, FLM viruses can be used to determine neutralizing antibodies in serum without the influence of weakly neutralizing FL- and prM-Abs (a better representation for type specific neutralizing potency)^35^ and as an antigen for serum depletion^36,37^. Importantly, our *in vitro* ADE assays clearly, and perhaps is the first time to directly demonstrate that FL-targeting antibodies are the main contributors to ADE in polyclonal serum. Therefore, generating FLM variants for all DENV serotypes will enable the targeted separation of weakly neutralizing antibodies from other antibody populations in complex polyclonal sera, providing an extra dimension for assessing serum quality and vaccine assessment.

In this study, we developed an nLuc-based ADE assay to improve throughput. As with other *in vitro* ADE assays, the magnitude of enhancement reported as fold-change (nLuc) or percentage of infected cells (flow cytometry) is inherently variable and can be influenced by multiple factors including expression level of CD32a on K562 cells (in human CD16a)^38^, baseline nLuc signal, and virus maturity or stability. To ensure data quality and plate-to-plate consistency, internal monoclonal antibody standards (4G2 and B7) were included on every assay plate. Although absolute enhancement values should not be overinterpreted, the ranges of antibody or serum concentrations that produce enhancement were highly consistent. Importantly, peak ADE concentrations should be evaluated alongside neutralization titers (EC_50_) to interpret the biological properties of a serum or antibody. Nevertheless, serum from D2-vFLM-immunized mice displayed clear heterotypic neutralization against DV4 but did not exhibit measurable ADE, indicating that these cross-reactive antibodies are non-enhancing. These antibodies may represent EDE-type responses or a distinct class of cross-reactive antibodies with minimal enhancement capacity. We are currently working to define the specificity of these antibodies.

Not all FL-targeting antibodies are weakly neutralizing, and some protective antibodies have overlapping footprints on the FL epitopes. Compared with other reported mutations in the FL region of flavivirus vaccine candidates, the N103S and G106L substitutions described here are relatively conservative, which may explain their retained functionality. This is important not only for LAV development but also for maintaining the overall immunogenicity of the vaccine. As shown in our neutralization data, both D2-vFLM and D4-vFLM remain sensitive to neutralization by EDE1-C10, a cross-reactive quaternary antibody with a limited footprint on the FL. Functionally preserving the FL may also help maintain the structural integrity of the FL region, thereby retaining epitopes recognized by other protective antibodies that partially overlap with the FL^39^.

Despite the negative perception surrounding the current DENV vaccines, all licensed formulations have been shown to be safe and protective in DENV-experienced populations, saving thousands of lives. Nevertheless, there remains room for further improvement in safety for DENV-naïve populations. Here, we report a targeted approach to enhance the safety of DENV-LAV. We also suggest that our FL mutations preserve the structure and function and may also have implications for other emerging vaccine platforms such as mRNA-LNP, E-dimer, and virus-like particle (VLP) vaccines. The transferability of the FLM modification to other DENV serotypes also enables rapid engineering of new strains and other orthoflaviviruses to address potential breakthrough infections caused by diverse DENV genotypes^40,41^. LAVs mimic natural infection, generating robust humoral and cell-mediated immune responses. They also represent the most practical and economical vaccine platform, capable of providing protection with a single dose in DENV endemic regions. To date, LAVs remain the only proven vaccine platform in clinical use with demonstrated efficacy against DENV infection. However, as history has shown, indiscriminately chasing immunogenicity of DENV vaccines can be risky especially when the ADE-prone FL-epitope is also immunodominant^14,42^. It is therefore our responsibility to ensure that future vaccine designs continue to balance both efficacy and safety for all populations.

## MATERIALS AND METHODS

### Cells and Viruses

*Aedes albopictus* C6/36 cells (ATCC CRL-1660) were cultured in MEM (Gibco) supplemented with 5% FBS (HyClone), 1% penicillin/streptomycin (Gibco), 0.1 mM nonessential amino acids (Gibco), 1% HEPES (Gibco), and 2 mM GlutaMAX (Gibco) at 32°C with 5% CO_2_. African green monkey Vero-81 cells (ATCC CCL-81) were maintained in DMEM/F12 (Gibco) supplemented with 10% FBS, 1% penicillin/streptomycin (Gibco), 0.1 mM nonessential amino acids (Gibco), and 1% HEPES (Gibco) at 37°C with 5% CO_2_. K562 cells were cultured in suspension in RPMI-1640 medium (Sigma-Aldrich) supplemented with 10% fetal calf serum (Cytiva), 2 mM L-glutamine, and 1% penicillin–streptomycin (Sigma-Aldrich) at 37°C in a humidified atmosphere containing 5% CO_2_. The medium was renewed every 2-3 days by adding or replacing fresh complete medium. DENV viruses (DV2-S16803, GenBank accession: GU289914; DV4-SriLanka-92, GenBank accession: KJ60504.1) were propagated in either C6/36 or Vero-81 cells using infection media. C6/36 infection media consisted of Opti-MEM (Gibco) supplemented with 2% FBS (HyClone), 1% penicillin/streptomycin (Gibco), 0.1 mM nonessential amino acids (Gibco), 1% HEPES (Gibco), and 2 mM GlutaMAX (Gibco). Vero-81 infection media consisted of DMEM/F12 (Gibco) supplemented with 2% FBS, 1% penicillin/streptomycin (Gibco), 0.1 mM nonessential amino acids (Gibco), and 1% HEPES (Gibco). D2-vFLM was adapted from previously described D2-FLM^19^ through 5 passages on Vero-81 at 32°C, followed by 5 passages on the same cell line at 37°C. Dengue nLuc reporter viruses were created using a four-plasmid system in which nLuc reporter was inserted into the viral genome as previously described^43^. In short, nLuc gene was cloned after the coding sequence of the first 38 amino acids of the DENV capsid gene, linked by the Thosea asigna virus 2A (T2A) self-cleaving peptide. Additionally, a codon-optimized C38 sequence was placed after T2A to prevent homologous recombination between the two C38 sequences. Stock viruses are deep-sequenced in-house using a iSeq100 platform with 500x - 1000x genome coverage.

### DENV focus-forming assay

Virus titers were determined using a focus-forming assay in a 96-well plate. Vero-81 and C6/36 cells were seeded at densities of 2×10^4^ and 1×10^5^ cells per well, respectively, one day before infection. The following day, virus was serially diluted 10-fold across eight dilutions, and 50 µL of each dilution was added to wells after removing the growth medium. The plate was incubated for 1 hour at 37°C (Vero-81) or 32°C (C6/36). Post-incubation, 125 µL of overlay medium (Opti-MEM supplemented with 2% FBS, 1% NEAA, 1% P/S, and 1% methylcellulose) was added to each well, and infection proceeded for 48 hours. The overlay was then removed, and wells were washed three times with PBS before fixation with 10% formalin in PBS for 30 minutes. Cells were blocked using a permeabilization buffer (ThermoFisher, 00-8333-56) containing 5% non-fat dried milk. Primary antibodies, anti-prM 2H2 and anti-E 4G2 (from non-purified hybridoma supernatant), were diluted 1:200 in blocking buffer, and 50 µL of the diluted primary antibody was added to each well. The plate was then incubated at 37°C for 1 hour. After incubation, wells were washed before adding 50 µL of a goat anti-mouse HRP secondary antibody (SeraCare KPL 54500011) diluted 1:2,000. The plate was incubated at room temperature for 1 hour, followed by additional washing steps. Foci were then stained using TrueBlue HRP substrate (SeraCare) and quantified with an ImmunoSpot analyzer (Cellular Technology).

### DENV infectivity assay using nLuc-reporter viruses

One day before infection, Vero-81 cells were seeded at 2×10^4^ cells per well in a black 96-well plate (Corning Costar). After 24 hours, the virus was serially diluted 10-fold across eight dilutions. The growth medium was removed from the cell plate, and 50 µL of each dilution was added to the corresponding wells. The plate was then incubated at 37°C for 48 hours. Following incubation, Nano-Glo Luciferase Assay substrate (Promega) was added to each well, following manufacturer’s protocol, and luciferase activity was detected using GloMax Plate Reader (Promega) as surrogates for viral infection.

### DENV maturation via western blot

Viral supernatants were mixed with 4x Laemmli Sample Buffer (Bio-Rad) and heated at 95°C for 5 minutes. Proteins were separated through SDS-PAGE electrophoresis, transferred to a PVDF membrane, and blocked with 5% nonfat milk in TBS-T. The membrane was incubated for 1 hour at 37°C with polyclonal rabbit anti-prM (1:1,000; Invitrogen PA5-34966) and polyclonal rabbit anti-Env (1:1,000; Invitrogen PA5-32246) in 2% BSA in PBS-T. After incubation, membrane was thoroughly washed with TBS-T and incubated for 1 hour at room temperature with goat anti-rabbit HRP (1:10,000; Jackson ImmunoLab) in 5% milk in TBS-T. Detection was performed using SuperSignal West Pico Plus chemiluminescent substrate (ThermoFisher), and imaging was carried out on ChemiDoc Imaging system (Bio-Rad). Band pixel intensity was quantified using Image Lab software (Bio-Rad), and relative maturation was determined using the equation: (prMExp/EnvExp)/(prMWT/EnvWT).

### DENV Growth Kinetics

Vero-81 and C6/36 cells were seeded on 6-well plates one day before infection, at a concentration of 4×10^5^ and 5×10^5^, respectively. Cells were infected with an MOI of 0.05-0.1, estimating 8×10^5^ (Vero-81) and 1×10^6^ (C6/36) cells on the day of infection. After virus was added to the cells, plates were incubated for 1 hour at 37°C (Vero-81) or 32°C (C6/36), followed by 3x washes with PBS and replenishment with fresh infection medium. At 0, 24, 48, 72, 96, and 120 hours after infection, 300 µL of viral supernatant was collected, followed by replenishing with the same amount of fresh infection media, and stored at −80°C. Dengue focus-forming assay was performed, as previously described, for each time point. All experiments were performed independently at least three times.

### Thermostability and pH stability assays

Viruses were thawed and incubated at 4°C, 32°C, 37°C, 42°C and 55°C for 1 hour, followed by virus quantification luciferase reporter assay, as previously described. All experiments were performed independently at least three times. For pH stability assay, we prepare infection medium adjusted at 4 different pH (5.4, 6.4, 7.4, and 8.4) using hydrochloric acid or sodium hydroxide. Viruses were mixed 1:1 with the pH-adjusted media and incubated for 1 hour at 4°C. Following incubation, virus quantification was performed using luciferase reporter assay, as previously described. All experiments were performed independently at least three times.

### Focus reduction and nLuc based-neutralization titer assays

Focus reduction neutralization titer (FRNT) assays were conducted as previously described. One day prior infection, 2×10^4^ Vero-81 cells were seeded in a 96-well plate. Antibodies were serially diluted and mixed with the DENV at 100 FFU/well (FRNT) or DENV-nLuc at 800 FFU/well (nLuc-based) at a 1:1 ratio and incubated for 1 hour. The mixture was then added to the cells and incubated for another hour before applying the overlay (Opti-MEM supplemented with 2% FBS, 1% NEAA, 1% P/S, and 1% methylcellulose). Plates were incubated for 48 hours, after which viral foci were stained and counted, as previously detailed in ‘DENV focus-forming assay’ or Nano-Glo Luciferase Assay substrate was added for the nLuc-based assay. The neutralization % is fitted to a variable slope sigmoidal dose-response curve, using GraphPad Prism 10. For serum neutralization, FRNT_50_ values were determined from an eight-point dilution curve starting at 1:40, followed by 4-fold serial dilutions. All experiments were performed independently at least three times.

### ADE assay using nLuc-reporter DENV

Antibody-dependent enhancement (ADE) was evaluated using K562 cells cultured in suspension. On day −1, K562 cells were seeded at >1×10^6^ cells/mL. On the day of experiment, cells were counted and seeded into black, flat-bottom 96-well plates at a density of 1×10^5^ cells per well. Ab or serum samples were serially diluted at four-fold interval for eight times in infection medium (RPMI-1640 supplemented with 2% fetal bovine serum and 1% penicillin–streptomycin), with starting dilution of 1:80. The virus was diluted in the same medium to achieve 5,000 focus-forming units (FFU) per well. Equal volumes of diluted serum and virus (25 µL each) were mixed and incubated at 37°C for 1h and added to the plated cells (50 µL per well), resulting in a total volume of 100 µL per well. Each plate included a negative control consisting of K562 cells infected with virus only, which served as a baseline for normalization and two mAb (4G2 and B7) standards. Plates were incubated for 24 hours at 37°C in a humidified 5% CO₂ incubator. Following incubation, the Nano-Glo Luciferase Assay substrate (Promega) was prepared according to the manufacturer’s instructions and added directly to each well. Luminescence was measured using a NanoGlo luciferase reader (Promega), and results were reported as fold-change compared to no Ab control.

### Next Generation Sequencing

Viral RNA was isolated using TRIzo LS (Invitrogen), following the manufacture’s protocol. Samples for next generation sequencing (NGS) library were generated via Click-seq protocol based on previously described work-flow^44^. In brief, reverse transcription was carried out using Superscript IV Reverse Transcriptase (Invitrogen) according to the manufacturer’s protocol, except that the reaction was supplemented with small amounts of azido-nucleotides (AzNTPs). ClickSeq was carried out for library preparation, incorporating Illumina TrueSeq Index Primers. Libraries were quantified using Quantus Fluorometer (Promega) and sequenced using Illumina iSeq 100 System with a pair-ended 150bp x 2 reagents. Each sample was aimed for 100x genome coverage with final coverage after QC ranging from 500x – 1000x. Raw fastq files were imported to Geneious Prime software for quality control and variant calling after aligned to the original viral genome.

### Mouse immunization

Male and female C57BL/6 (B6) mice, 3-4 months of age, were used for immunizations. Animals were randomly assigned to receive either the DV2 or D2-vFLM viruses, with each group consisting of 10 - 11 animals. Each mice received 2.5×10^5^ FFU DENV2 in PBS and mixed with an equal volume of AddaVax adjuvant (InvivoGen). Immunizations were delivered intramuscularly into the hind leg. The schedule consisted of a prime immunization, followed by a first boost two weeks later and a second boost two weeks after the first boost. Blood samples were collected prior to each immunization by submandibular bleed using BD Microtainer SST gel tubes. Two weeks after the second boost, mice were euthanized, and blood was obtained by cardiac puncture. Samples were allowed to clot at room temperature and serum were collected after low-speed centrifugation. The resulting serum was transferred to 1.5 mL microcentrifuge tubes and heat-inactivated at 55 °C for 30 minutes before storage and further analysis.

### AG129 challenge model

Male and female AG129 mice, 4-6 months-old were immunized with 5×10^4^ FFU of DV2, D2-vFLM or mock with AddaVax (3:1) immunized with the same route and schedule as described above. Three weeks after the final boost, mice were infected with 5×10^5^ FFU of mouse-adapted D220 via tail-vein intravenous (I.V.) injection in PBS. Body weights were measured daily for 10 days. Serum was collected via cheek bleed as described above to monitor viral titer from d1pi to d4pi. Mice whose body weight dropped below 80% of their initial weight were humanely euthanized using CO_2_ overdose followed by cervical dislocation as a secondary confirmatory method. All animal experiments were approved by the Saint Louis University Institutional Animal Care and Use Committee (IACUC).

### Statistical analysis

GraphPad Prism version 9.0 was used for statistical analysis. Titer and % infection of D2-vFLM were compared to the DV2 using two-way ANOVA. EC_50_ and FRNT_50_ were compared using Student’s t-test. Significant symbols are as follows: *, P < 0.05; **, P< 0.005; ***, P < 0.0005; ****, P < 0.00005. The data are graphed as means ± standard deviations. All experiments were conducted at least three times, except for experiments involving serum were performed at least 2 times due to limited regents.

## ACKNOWLEDGEMENTS

We thank members of the Tse laboratory for helpful discussions. We also thank Department of Comparative Medicine at Saint Louis University for providing guidance and help for animal experiments. We also thank Dr. Eva Harris at UC Berkeley for providing the D220 mouse adaptive DENV2 challenge strain. This work was supported by Saint Louis University Department of Molecular Microbiology and Immunology Startup fund, SLU-SOM President’s Research Award, and Institute for Drug and Biotherapeutic Innovation Seed Grant.

## AUTHOR CONTRIBUTION

L.V.T. designed the study. Y.R.D, J.J.A., K.D., L.J.S., and L.V.T. performed experiments. L.V.T. and R.M.M. performed high-throughput sequencing preparation and analysis. provided reagents. L.V.T. provided oversight of the project and funding. L.V.T. wrote the manuscript. Y.R.D., R.M.M. and L.V.T. reviewed and revised the final version. All authors approved the final version of the manuscript.

---

**Figure S1.**
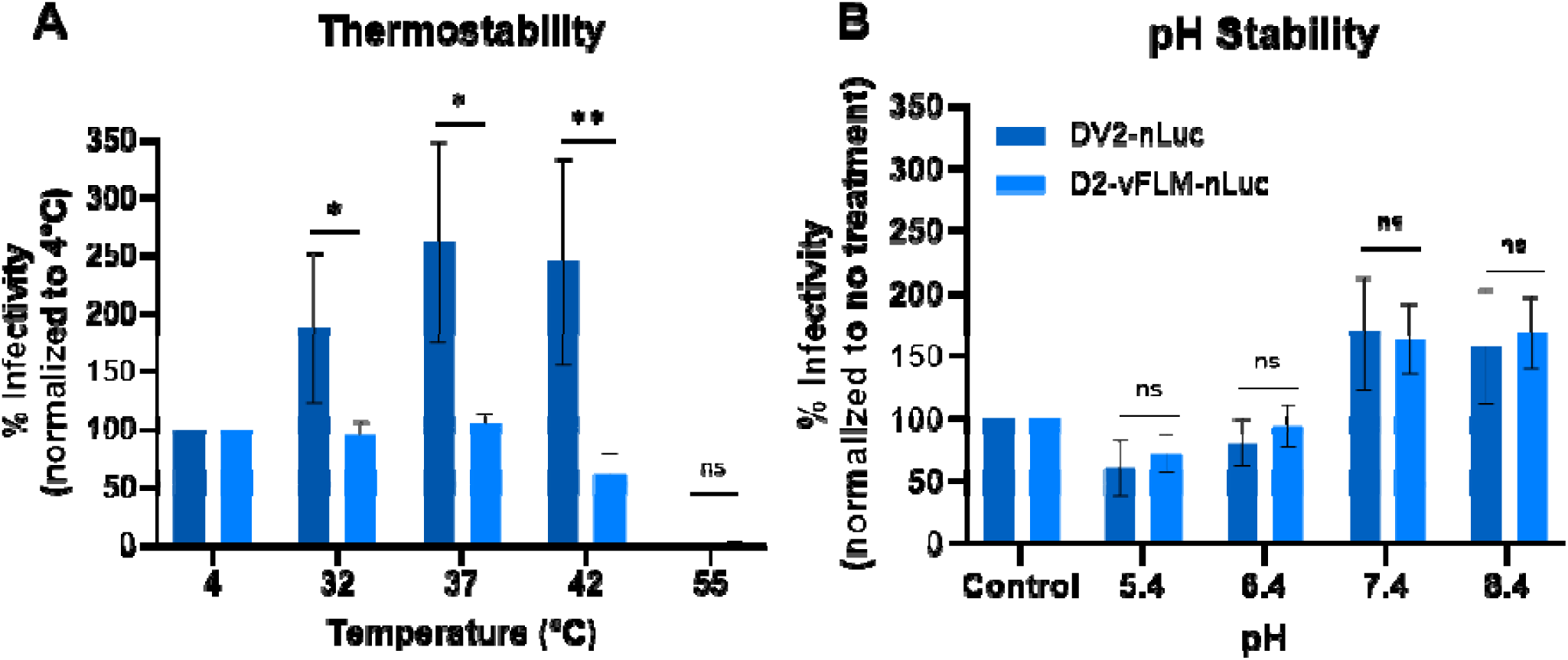
Temperature and pH stability of DV2 and D2-vFLM assessed using a nLuc reporter assay. **A)** Infectivity of DV2-nLuc and D2-vFLM-nLuc following incubation at temperatures ranging from 4°C to 55°C, normalized to the control 4°C condition. **B)** Infectivity of DV2-nLuc and D2-vFLM-nLuc following exposure to buffers at pH 8.4, 7.4, 6.4, or 5.4, normalized to the untreated control. Statistics were performed by Students *t-*test.

**Figure. S2.**
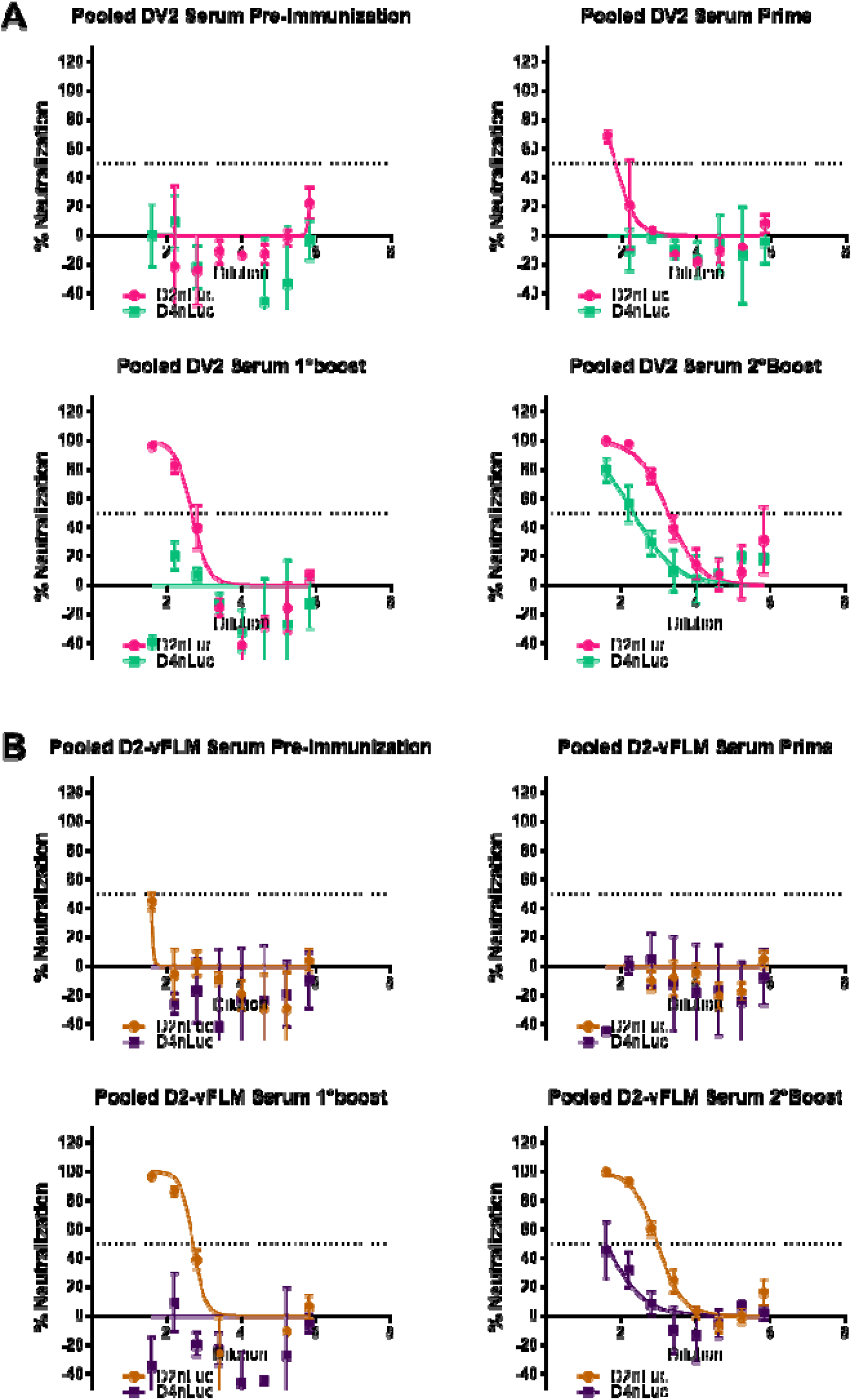
Neutralization curves of pooled serum samples from **A)** DV2 and **B)** D2-vFLM immunized mice against homotypic and heterotypic DENV-nLuc.

**Figure S3.**
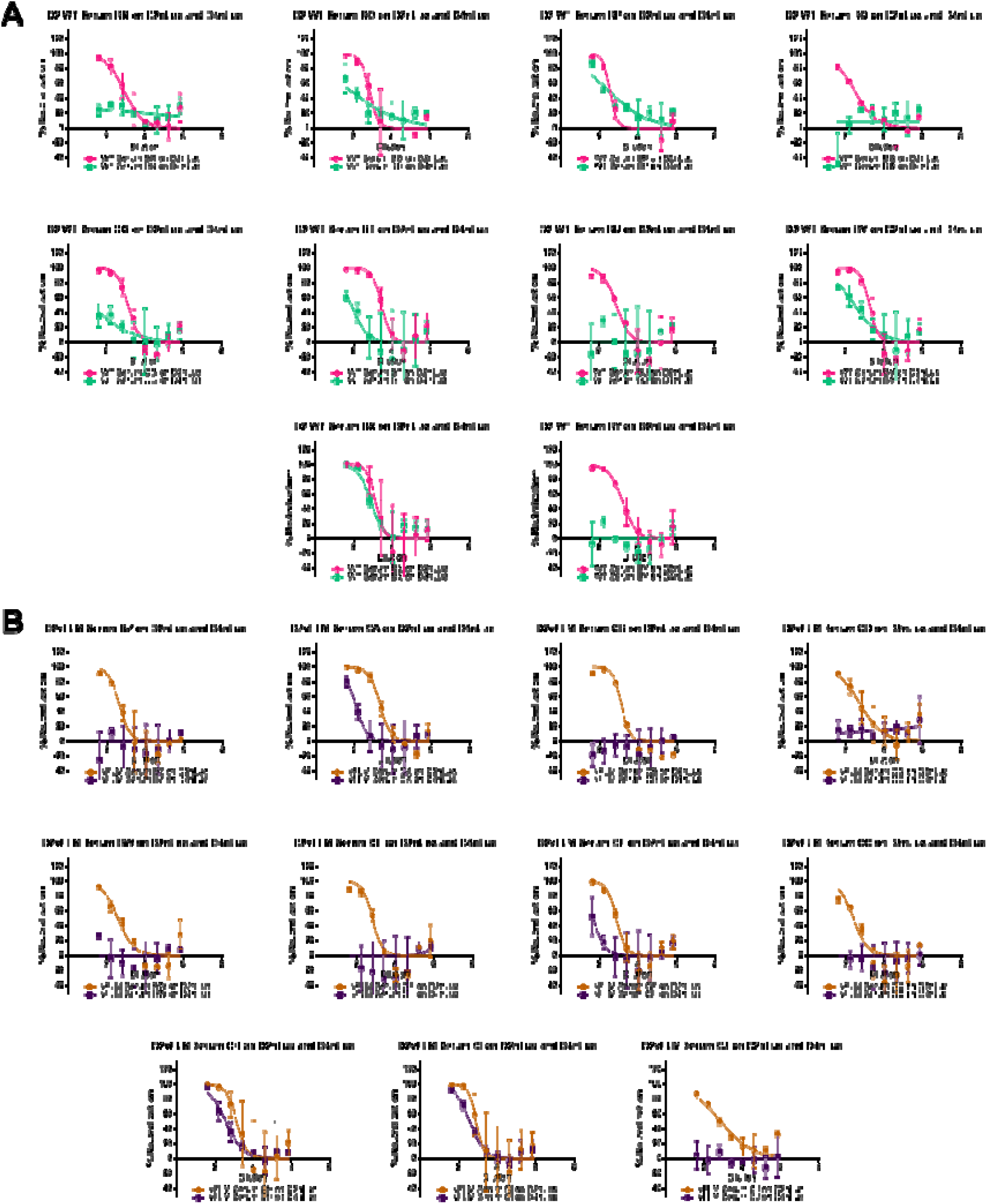
Individual neutralization curves of **A)** DV2 and **B)** D2-vFLM serum after 2°boost against homotypic (DV2-nLuc) and heterotypic (DV4-nLuc).

**Figure S4.**
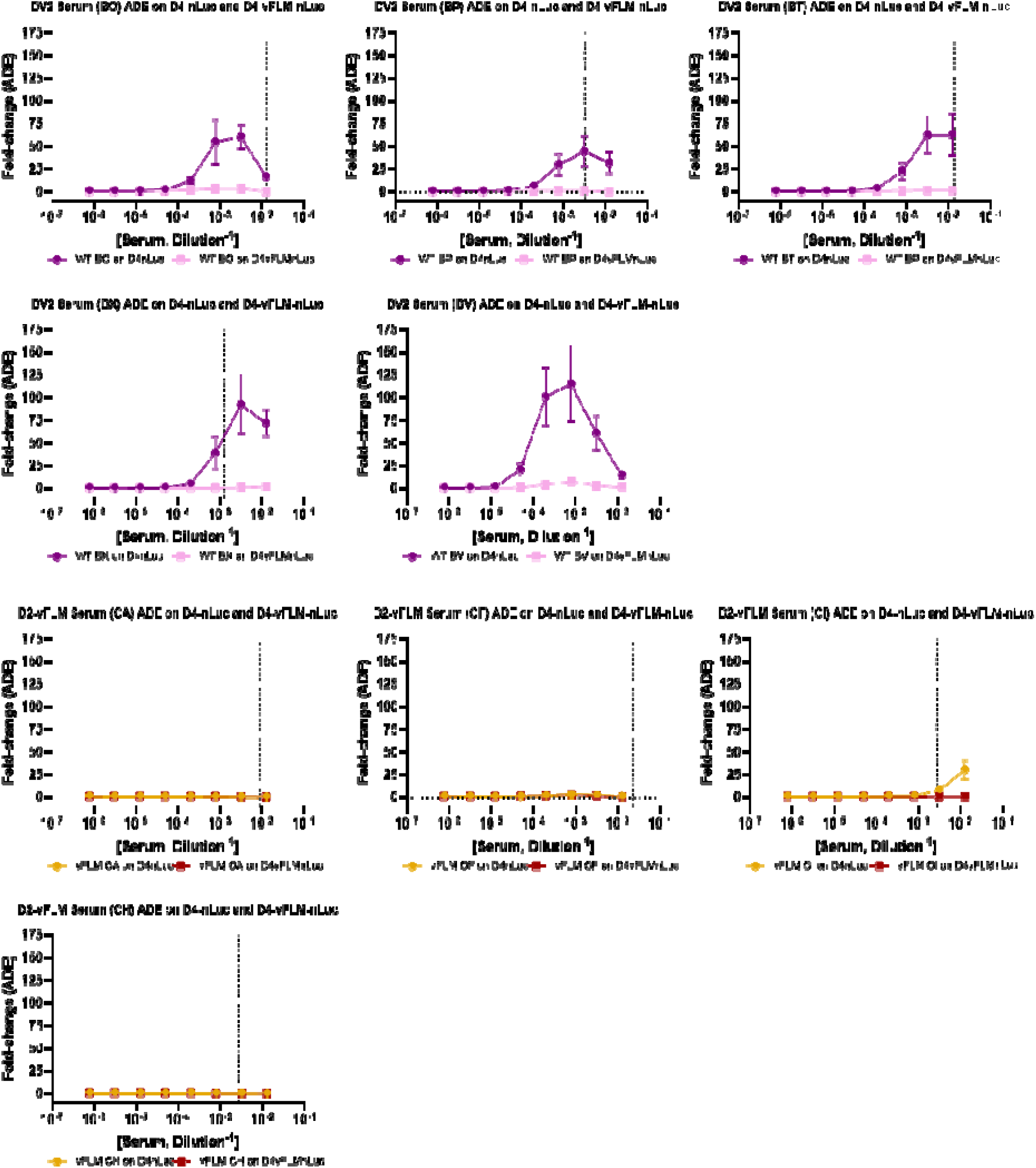
Individual heterotypic ADE curves of DV4-nLuc and D4-vFLM-nLuc against DV2 and D2-vFLM serum from immunized mice. Vertical dotted lines denote the EC50 of the serum against DV4-nLuc.

